# A metric on phylogenetic tree shapes

**DOI:** 10.1101/054544

**Authors:** C. Colijn, G. Plazzotta

## Abstract

The shapes of evolutionary trees are influenced by the nature of the evolutionary process, but comparisons of trees from different processes are hindered by the challenge of completely describing tree shape. We present a full characterization of the shapes of rooted branching trees in a form that lends itself to natural tree comparisons. The resulting metric distinguishes trees from random models known to produce different tree shapes. It separates trees derived from tropical vs USA influenza A sequences, which reflect the differing epidemiology of tropical and seasonal flu. We extend the shape metric to incorporate summary features such as asymmetry, or statistics on branch lengths. Our approach allows us to construct addition and multiplication on trees, and to create a convex metric on tree shapes which formally allows computation of average trees.

## 1 Introduction

The availability and declining cost of DNA sequencing mean that data on the diversity, variation and evolution of organisms is more widely available than ever before. Increasingly, thousands of organisms are being sequenced at the whole-genome scale (Chewapreecha et al., 2014; Bedford et al., 2015; Anopheles gambiae 1000 Genomes, 2016). This has had particular impact on the study of pathogens, whose evolution occurs rapidly enough to be observed over relatively short periods. As the numbers of sequences gathered annually grow to the tens of thousands in many organisms, comparing this year’s evolutionary and diversity patterns to previous years’, and comparing one location to another, has become increasingly challenging. Despite the fact that evolution does not always occur in a tree-like way due to the horizontal movements of genes, phylogenetic trees remain a central tool with which we interpret these data.

The shapes of phylogenetic trees are of long-standing interest in both mathematics and evolution (Stam, 2002; Slowinski, 1990; Guyer & Slowinski, 1993; Purvis, Fritz, Rodríguez, Harvey, & Grenyer, 2011; Kirkpatrick & Slatkin, 1993; Mooers & Heard, 1997; Blum & François, 2006; Wu & Choi, 2015). A tree’s shape refers to the tree’s connectivity structure, without reference to the lengths of its branches. A key early observation was that trees reconstructed from evolutionary data are more asymmetric than simple models predict (Aldous, 1996). This spurred an interest in ways to measure tree asymmetry (Kirkpatrick & Slatkin, 1993; Fusco & Cronk, 1995; DJ, 2001; Pompei, Loreto, & Tria, 2012; Stich & Manrubia, 2009), in the power of asymmetry measures to distinguish between random models (Agapow & Purvis, 2002; Kirkpatrick & Slatkin, 1993; Matsen, 2006), and in constructing generative models of evolution that produce imbalanced trees (DJ, 2001; Manceau, A, & Morlon, 2015; Blum & François, 2006). Tree shapes carry information about the underlying evolutionary processes, and distributions of tree shapes under simple null models can be used to test hypotheses about evolution (Mooers & Heard, 1997; Blum & François, 2006; Blum, François, & Janson, 2006; Purvis et al., 2011; Wu & Choi, 2015). Recent work also relates fitness, selection and a variety of ecological processes to tree shape (LP, Colato, & Fontanari, 2004; Dayarian & Shraiman, 2014; J Hein, 2004; Wakeley & Wakeley, 2009; Gascuel, 2000; Manceau et al., 2015). An additional motivation for studying the shapes of phylogenetic trees is that reconstructing branch lengths is challenging, particularly deep in a tree. There may be weak support for a molecular clock, and coalescent inference procedures may produce trees with consistent shape but differing root heights.

Tree shape is well established as carrying important information about macroevolutionary processes, but also carries information about evolution in the short term. In the context of pathogens, diversity patterns represent a combination of neutral variation that has not yet become fixed, variation that is under selection, complex demographic processes (host behaviour and contact patterns), and an array of ecological interactions. The extent to which tree shapes are informative of these processes is not well understood, though there have been studies on the frequency of cherries and tree imbalance (Volz, Koelle, & Bedford, 2013; Lambert & Stadler, 2013; Plazzotta, Kwan, Boyd, & Colijn, 2016) and simulation studies aiming to explore the question (Leventhal et al., 2012; K. Robinson, Cohen, & Colijn, 2012; Colijn & Gardy, 2014; Plazzotta & Colijn, 2016).

A key limitation in relating tree shapes to evolution and ecology has been the limited tools with which trees can be quantified and compared. Comparing tree shapes from different models of evolution or from different datasets requires comparing *unlabelled* trees, whereas most tree comparison methods (eg the Robinson-Foulds (D. Robinson & Foulds, 1981), Billera-Holmes-Vogtmann (Billera, Holmes, & Vogtmann, 2001) and newer metrics (Kendall & Colijn, 2016)) compare trees with one particular set of organisms at the tips (one set of taxa, with the labels in each tree). These metrics can be used as a basis for metrics on unlabelled shapes, for example by setting the distance between shapes *T*_1_ and *T*_2_ to be 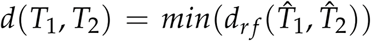, where 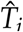 has shape *T_i_* and the Robinson Foulds distance is computed by labelling 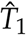 and 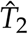 with the same set of labels. However, this requires computing the distance using every distinct arrangement of tip labels on one of the trees. Similarly defined metrics on trees with multi-sets for their labels have been described (Huber, Spillner, Suchecki, & Moulton, 2011), but their computation is difficult and metrics may not be applicable if the trees have different numbers of tips. Consequently, the tools at our disposal to describe and compare tree shapes from *different* sets of tips are limited, and have focused on scalar measures of overall asymmetry (Slowinski, 1990; Guyer & Slowinski, 1991; Matsen, 2006; Pompei et al., 2012; Fusco & Cronk, 1995; Stich & Manrubia, 2009; Colless, 1995; Sackin, 1972) and on the frequencies of small subtree shapes such as cherries (Steel & McKenzie, 2000; Plazzotta & Colijn, 2016; Volz et al., 2013) and r-pronged nodes (Rosenberg, 2006). Recently, kernel (Poon et al., 2013) and spectral (Lewitus & Morlon, 2015) approaches also have been used.

Here we give a simple characterization of all possible shapes for a rooted tree and use this to define metrics (in the sense of true distance functions) on tree shapes. The scheme provides an efficient way to count the frequencies of sub-trees in large trees, and hence can be used to compare empirical distributions of sub-tree shapes. It is not limited to binary trees and can be formulated for any maximum size multifurcation, as well as for trees with internal nodes with only one descendant (sampled ancestors). As an illustrative example, we apply a metric derived from our scheme to simulated and data-derived trees. It separates trees from random tree models that are known to produce trees with different shape. We use the approach to compare trees from tropical vs USA human influenza A (H3N2). We extend the metric to incorporate statistics on the lengths of branches or other tree features, and we use a map from tree shapes to the rational numbers to define a convex metric on tree shapes.

## 2 Materials and Methods

### 2.1 Definitions

A *tree shape* is a tree (a graph with no cycles), without the additional information of tip labels and branch lengths. We consider rooted trees, in which there is one node specified to be the root. Tips, or leaves, are those nodes with degree 1. A *rooted tree shape* is a tree shape with a vertex designated to be the root. We use "tree shape", as we assume rootedness throughout. Typically, edges are implicitly understood to be directed away from the root. A node’s *children* are the node’s neighbours along edges away from the root. A *multifurcation*, or a *polytomy*, is a node with more than two children, and its *size* is its number of children (> 2). Naturally, a rooted phylogeny defines a (rooted) tree shape if the tip labels and edge weights are discarded. Phylogenies typically do not contain internal nodes with fewer than two children (sampled ancestors), but we allow this possibility in the tree shapes.

### 2.2 Labelling scheme

Our approach is to label any possible tree shape, traversing the tree from the tips to the root and assigning labels as we go. The simplest case is to assume a binary tree, in which all internal nodes have two children. We give a tip the label 1. For every internal node, we list its childrens’ labels (*k*, *j*). Using lexicographic sorting, list all possible labels (*k*, *j*): (1), (1, 1), (2, 1), (2, 2), (3, 1), (3, 2), (3, 3),… We define the label of a tree shape whose root node has children (*K*, *J*) to be the index at which (*K*, *J*) appears in this list. Accordingly, a “cherry” (a node with two tip children) is labelled 2 because its children are (1, 1), which is the second entry in the list. A node with a cherry child and a tip child (a (2, 1), or a pitchfork) has label 3. The tree shape (*k*, *j*) (a tree whose root has a child with label *k* and one with label *j*) has label 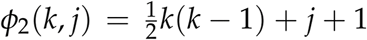. To see this, note that there are *k* pairs of the form (*k*, *j*) (with 1 ≤ *j* ≤ *k*). We label tips as 1, and the first pair (1, 1) has label 2. Sorting lexicographically, this means that the pair (*n*, 1) has label 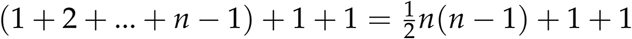, because all labels starting with 1, 2, 3,…*k*,..*n* − 1 occur before this first pair beginning with *n*. The extra 1 accounts for starting the scheme with (1, 1) whose label is 2. Then (*k*, *j*) has label 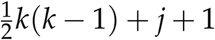. We continue until the root of the tree has a label.

Each node in a tree is therefore labelled based on its childrens’ labels. If the trees are full binary trees (the simplest case), we call the label function *ϕ*_2_:

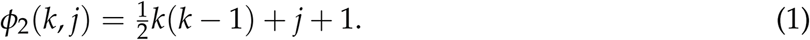

The subscript 2 specifies that each node has a maximum of two children; the scheme can be extended but has a different explicit form (*ϕ_M_*) if there are multifurcations or internal nodes with a single child (in which case we require *j* ≥ 0 rather than 1). We give details in the Supplementary Materials.

### 2.3 Metrics on the space of rooted unlabelled shapes

This characterisation leads to simple metrics on the space of tree shapes. The simplest is a comparison of the root labels: given two binary trees *T_a_* and *T_b_*, whose root nodes are *R_a_* and *R_b_*, and where the label of a node *x* is *L*(*x*), we can write

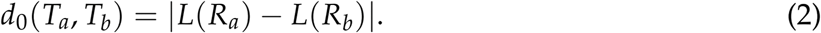

In other words, the absolute difference between the root nodes’ labels is a metric, with tree 1 a distance of 1 from tree 2 and so on. Clearly *d*_0_ is symmetric and non-negative. The tree isomorphism algorithm and the above labelling clearly show that 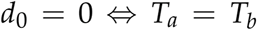 and the absolute value obeys the triangle inequality. However, *d*_0_ is not a very useful metric, in the sense that a large change in root label can result from a relatively “small” change in the tree shape (such as the addition of a tip).

While each tree is defined by the label of its root, it is also defined (redundantly) by the labels of all its nodes. If the tree has *n* tips, the list of its labels contains *n* 1s, typically multiple 2s (cherries) and so on. Let *L_a_* denote the list of labels for a tree 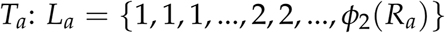. The label lists are multisets because labels can occur multiple times. Define the distance *d*_1_ between *T_a_* and *T_b_* to be the number of elements in the symmetric set difference between the label lists of two trees:

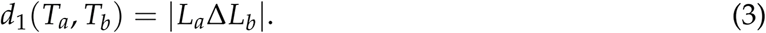

The symmetric set difference 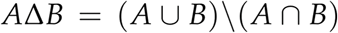 is the union of *A* and *B* without their intersection. Intuitively, this is the number of labels not included in the intersection of the trees’ label lists. If *A* and *B* are multisets with *A* containing *k* copies of element *x* and *B* containing *m* copies of *x*, with *k > m*, we consider 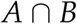 to contain *m* copies of *x* (these are common to both *A* and *B*). *A*∆*B* has the remaining *k* – *m* copies. Each tree’s label list contains more 1s (tips) than any other label. Accordingly, this metric is most appropriate for trees of the same size, because if trees vary in size, the metric can be dominated by differences in the numbers of tips. For example, if *L_a_*= {1, 1, 1, 1, 2, 2} (four tips joined in two cherries) and *L_b_*= {1, 1, 1, 2, 3} (three tips, i.e. a pitchfork), then 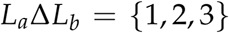, because there is a 1 and a 2 in *L_a_* in excess of those in *L_b_*, and a 3 in *L_b_* that is not matched in *L_a_*. Like *d*_0_, *d*_1_ is a metric: positivity and symmetry are clear from the definition. The cardinality of the symmetric difference is 0 if and only if the two sets are the same, in which case the root label is the same and the tree shapes are the same. That the symmetric difference obeys the triangle inequality is readily seen from the property 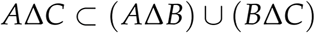.

Perhaps the most natural metric based on the labels, and the metric that we apply (and extend) through this work, compares the numbers of occurrences of each label in each tree. Let *v_a_* be a vector whose *k'* th element *v_a_*(*k*) is the number of times label *k* occurs in the tree *T_a_*; so *v_a_*(1) will be the number of tips, *v_a_*(2) the number of cherries, and so on. Define the metric

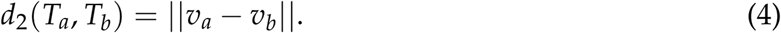

Positivity, symmetry and the triangle inequality are evident, and again *d*_2_ can only be 0 if *T_a_* and *T_b_* have the same number of copies of all labels (including the root label), which is true if and only if *T_a_* and *T_b_* have the same shape. This has a similar flavour to the statistic used to compare trees to Yule trees in (Blum & François, 2006), where the numbers of clades of a specific size were compared. We have used and extended metric *d*_2_ in the analyses presented in the Results.

Each of these metrics is computed in linear time. If *T_a_*, *T_b_* have *n_a_*, *n_b_* internal nodes, computing the distance requires 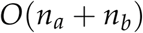 operations to define the labels, and 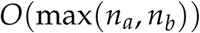 operations to compare the lists of labels. Different choices of weights increase computational time but not computational complexity; the variants we present are all linear in the (maximum) number of tips of the two trees.

### 2.4 Addition and multiplication of tree shapes defined by the mapping

Natural metrics associated with the labelling scheme are all based on the bijective map *ϕ* between the tree space 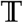 and the natural numbers 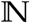. Composing *ϕ* with bijective maps between 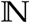 and other countable sets like the integers 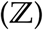, the positive rational numbers 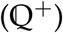, or the rationals 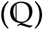 opens up further possibilities because we can take advantage of the properties (addition, multiplication, distance, etc) of integer and rational numbers. If *ψ* is a bijective map between 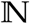 and one of these sets, then the composition 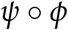 is also bijective, and we can use it to define addition and multiplication operations on trees:

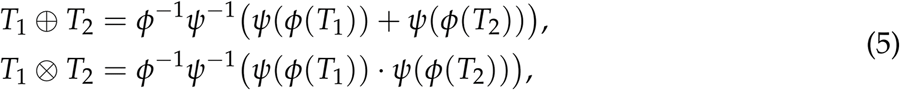

where + and · are the usual addition and multiplication. Now the space of trees together with these definitions of addition and multiplication, 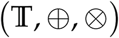, inherits all the algebraic properties of the set it is mapped into. For instance, 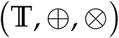 is a commutative ring if 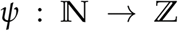. These constructions allow algebraic operations in the tree space 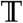. However the choice of the map *ψ* determines whether these operations are “meaningful” or “helpful” for applications of branching trees in biology or other fields. It turns out that the selection of a meaningful map is challenging.

For example, we can use the labelling scheme to map tree shapes to the (positive and negative) integers. We first extend *ϕ* with 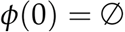, i.e. the the empty tree no tips. Consider the following well-known map between 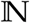 and 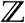:

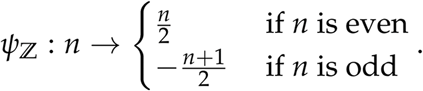

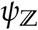 is clearly bijective: each tree shape is mapped to a unique integer and each integer corresponds to a unique tree shape. A representation of ten trees is provided in Figure S5. To "add" or "multiply" trees, we can add or multiply their corresponding integers and then invert, as in Eq (5). This may seem intuitive for small trees; for example the sum of tree number 3 and tree number -1 gives tree number 2 which has one fewer tip than tree number 3. For larger trees, however, addition and multiplication operations are less intuitive and do not follow the numbers of tips. This map has the advantage of simplicity but results in a large distance between trees differing by one tip.

### 2.5 Mapping tree shapes to the rational numbers

We use the map to the integers, and a map to the rational numbers, to define a *convex* metric on tree shapes. Convexity is the property that there is a point directly in between two other points (so that equality holds in the triangle inequality). Define the following map from 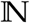 to 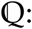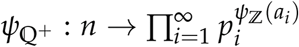 if *n >* 0, or 0 if *n* = 0. Here, *p_i_* are all the prime numbers and 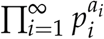 is the unique prime decomposition of *n* + 1. 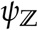 is as defined above, mapping the positive integers to all integers. For example 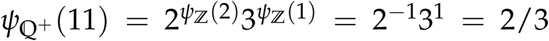. 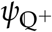 is injective, from the uniqueness of the prime factorization and the injectivity of because 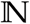 and 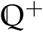 have the same cardinality. Therefore 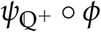 maps tree shapes bijectively to the non-negative rational numbers. In turn, 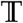 inherits all of the properties and structure of 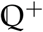. A distance metric 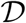 on 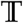 can be defined from the usual distance *| · |* of 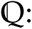

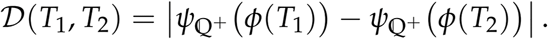

Because the absolute value is a convex metric in 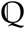, this is a convex metric on unlabelled tree shapes. It can be used to find averages of a set of trees. Figure S5 illustrates tree shapes together with their labels under the map 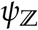.

### 2.6 Simulations

We compared trees from different random processes and models. One of the most natural random processes modelling phylogenetic trees is the continuous-time homogeneous birth-death branching process, in which each individual gives rise to a child at a constant rate in time, and also risks removal (death) at a constant rate. With birth rate *λ* and death rate *µ*, the ratio *λ*/*µ* specifies the mean number of offspring of each individual in this process, and affects the shapes and branching times of the resulting branching trees. In the epidemiological setting, the link to branching times has been used to infer the basic reproduction number *R*_0_ from sequence data (Stadler et al., 2012; Stadler, Kühnert, Rasmussen, & du Plessis, 2014). We computed the distances between trees derived from constant-rate birth-death (BD) processes simulated in the package `TreeSim` in R (Stadler, 2010). One challenge is that the number of tips in the BD process after fixed time is highly variable and depends on *λ*/*µ*. We aimed to detect shape differences that were not dominated by differences in the number of tips. Accordingly, we conditioned the processes to have 1500 taxa and then pruned tips uniformly at random to leave approximately 1250 tips remaining.

There are several other random models for trees. The Yule model is a model of growing trees in which lineages divide but do not die; in terms of tree shape it is the same as the Kingman coalescent and the equal rates Markov models. In the ‘proportional to distinguishable arrangements’ (PDA) model, each unlabelled shape is sampled with probability proportional to the number of *labelled* trees on *n* tips with that unlabelled shape (Rosen, 1978; Mooers & Heard, 1997). The “biased” model is a growing tree model in which a lineage with speciation rate *r* has child lineages with speciation rates *pr* and (1 – *p*)*r*. The Aldous’ branching model that we use here is Aldous’ *β*-splitting model with *β* = −1 (Aldous, 1996); in this model a *β* distribution determines the (in general asymmetric) splitting densities upon branching. The Yule, PDA, biased and Aldous *β* = −1 models are available in the package `apTreeshape` in R (Bortolussi, Durand, Blum, & O, 2006). We used *p* = 0.3 for the biased model, and sampled trees with 500 tips.

### 2.7 Data

We aligned data of HA protein sequences from human influenza A (H3N2) in different settings reflecting different epidemiology. Data were downloaded from NCBI on 22 Jan 2016. In all cases we included only full-length HA sequences for which a collection date was available. The USA dataset (*n* = 2168) included USA sequences collected between March 2010 and Sept 2015. The tropical data (*n* = 1388) included sequences from the tropics collected between January 2000 and October 2015. Accession numbers are included in the Supporting Information. Fasta files were aligned with mafft. Within each dataset, we sampled 500 taxa uniformly at random (repeating 200 times) and inferred a phylogenetic tree with the program FastTree. Where necessary we re-aligned the 500 sequences before tree inference. This resulted in 200 trees, each with 500 tips from the tropical and USA isolates.

Note that this approach is distinct from Bayesian inference of many trees on *one* set of tips, and from bootstrap trees on one set of tips. Either a posterior or bootstrap collection of trees from the same set of tips will share shape features because of the phylogenetic signal in the data. In contrast, we resample from the isolate collection each time and the trees we compare do not have the same set of labels.

### 2.8 Implementation

We have used R throughout. An R package is available on github at https://github.com/carolinecolijn/treetop. The implementation assumes full binary trees and includes metrics *d*_1_ and *d*_2_ with the option of weighting, as well as a “tree lookup” function that returns the tree associated with an integer in labelling scheme *ϕ*_2_.

## 3 Results

### 3.1 Label-based description of tree shapes

Figure 1 illustrates the labels at the nodes of two binary trees. The label of the root node uniquely defines the tree shape. Indeed, tree isomorphism algorithms use similar labelling (Lueker & Booth, 1979; Hopcroft & Tarjan, 1972; W, 1979; Colbourn & Booth, 1981; Sayward, 1981). If *R_a_* and *R_b_*are the root nodes of binary trees *T_a_* and *T_b_*, the tree shapes are the same if and only if 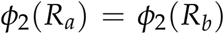. The map between trees and labels is bijective: every positive integer corresponds to a unique tree shape and vice versa.

Metrics are an appealing way to compare sets of objects; defining a metric defines a *space* for the set of objects – in principle allowing navigation through the space, study of the space’s dimension and structure, and the certainty that two objects occupy the same location if and only if they are identical. The labelling scheme gives rise to several natural metrics on tree shapes, based on the intuition that tree shapes are similar when they share many subtrees with the same labels.

### 3.2 Simulated random trees

There are several ways to sample random trees in ways known to produce trees of different shapes (in particular, different asymmetry). These include models capturing equal vs different speciation rates, continuous time birth-death processes with different rates and others (see Methods). We used the metric arising from our labelling scheme to compare these. Figure 2 shows a visualization of the tree-tree distances between trees from different random models. The metric groups trees from each process together and distinguishes between them well. Summary statistics such as tree imbalance also distinguish some of these groups well (particularly the PDA, Aldous, Yule and biased speciation model); indeed, we have elsewhere related the basic reproduction number to the number of cherries (Plazzotta & Colijn, 2016), and since the cherry is a symmetric configuration, trees with a high frequency of cherries will be more symmetric than those with a low frequency of cherries.

**Figure 1:**
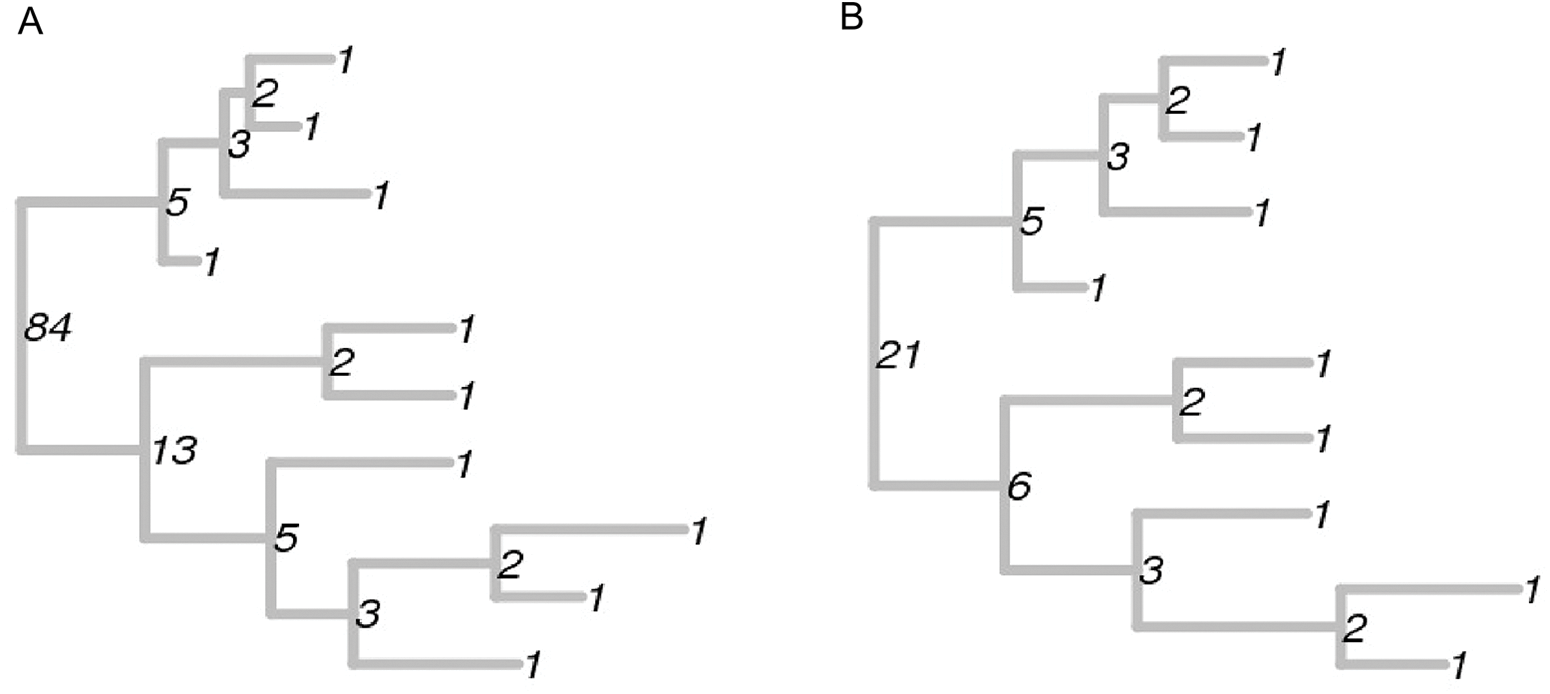
Illustration of the labels of the nodes of binary trees. Tips have the label 1. Labels of internal nodes are shown in black. The only difference between the trees in (a) and (b) is that in (b), the bottom-most tip from (a) has been removed. As a consequence, most of the labels are the same.

### 3.3 Tropical vs seasonal influenza

We also compared trees inferred from sequences of the HA protein in influenza A H3N2 sequences. Influenza A is highly seasonal outside the tropics (Russell et al., 2008), with the majority of cases occurring in winter. In contrast, there is little seasonal variation in transmission in the tropics. In addition, over long periods of time, influenza evolves in response to pressure from the human immune system, undergoing evolution particularly in the surface HA protein. This drives the ‘ladder-like’ shape of long-term influenza phylogenies (Koelle, Khatri, Kamradt, & Kepler, 2010; Volz et al., 2013; Westgeest et al., 2012; Luksza & Lässig, 2014), but would not typically be apparent in shorter-term datasets. With this motivation, we compared tropical samples to USA sample. Figure 3 shows that the tropical and USA flu trees are well separated by the metric. In addition, we used discriminant analysis of principal components (DAPC) (Jombart, Devillard, & Balloux, 2010) to determine which shapes separate the two groups. These shapes are those with high loadings on the first (and only substantial) principal component. We show them in Figure 3, listing their labels and colouring them according to Sackin imbalance. The two groups are different in imbalance, and the metric allows us to determine which sub-shapes occur with different frequencies to separate the groups. In the Supplementary Materials we compare the imbalance and numbers of cherries across the various groups of trees.

**Figure 2:**
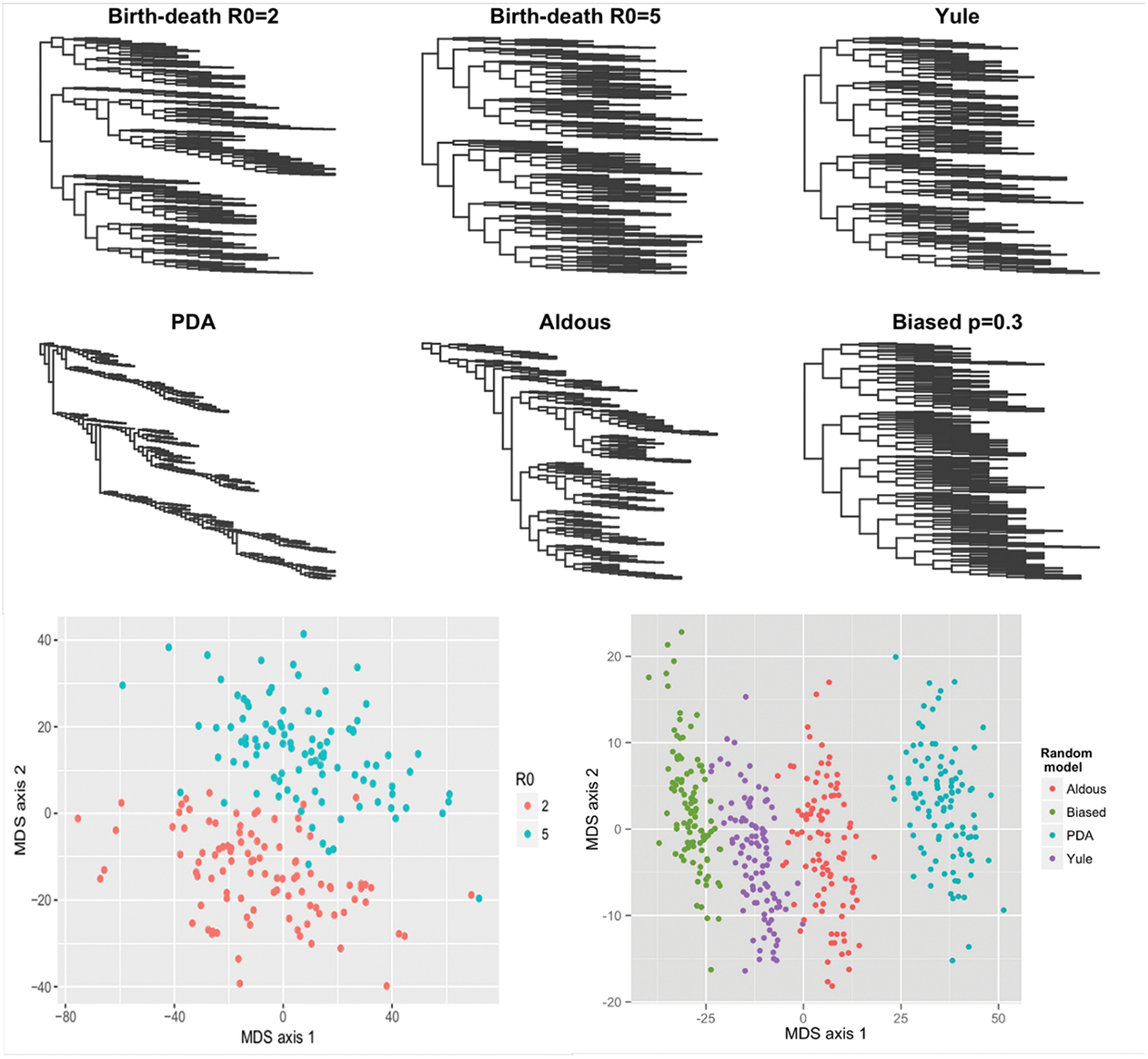
Top: Six sample trees, one from each of six different random processes. Bottom: Multi-dimensional scaling (MDS) plots showing that trees from each process are grouped together in the metric. Bottom left: trees from a birth-death model with different values of *R*_0_ = *λ*/*µ*. Bottom right: trees derived from the Yule, proportional to distinguishable arrangements (PDA), Aldous and biased models, each with 500 tips.

### 3.4 Incorporating tree size, branch lengths and other properties

Perhaps as it should be, the dominant difference between a tree with ten tips and one with one hundred tips is the size of the tree (and for this reason we have focused our application on comparing trees of the same size). The largest contribution to the distances will result from comparing the number of instances of the label 1 (tip) in two trees; this is necessarily larger than any other label copy number, and furthermore, a tree with more tips can have more cherries, pitchforks and any other subtree than a tree with fewer tips.

However, it is straightforward to construct metrics that compare tree shapes of different sizes with respect to their proportional frequencies of sub-trees. We based the metric *d*_2_ on vectors whose *i*^th^ components were the number of sub-trees of label *i*; we can divide these vectors by the number of tips *n_a_* in the tree: 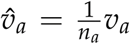, and include a component of 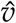 proportional to the number of tips ie 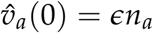. We then define a new Euclidean metric

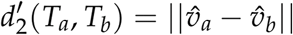

with *ɛ* > 0. With small *ɛ*, 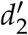 will be small when the proportional frequencies of sub-trees are very similar (even if the trees are different sizes), but will only be 0 if the trees have identical vectors and the same number of tips. Furthermore, if there are particular labels *i* that are of interest - for example those with relatively few tips, for a "tip-centric" tree comparison, weights *w* can be chosen and applied to the vectors to emphasize some entries more than others:

**Figure 3:**
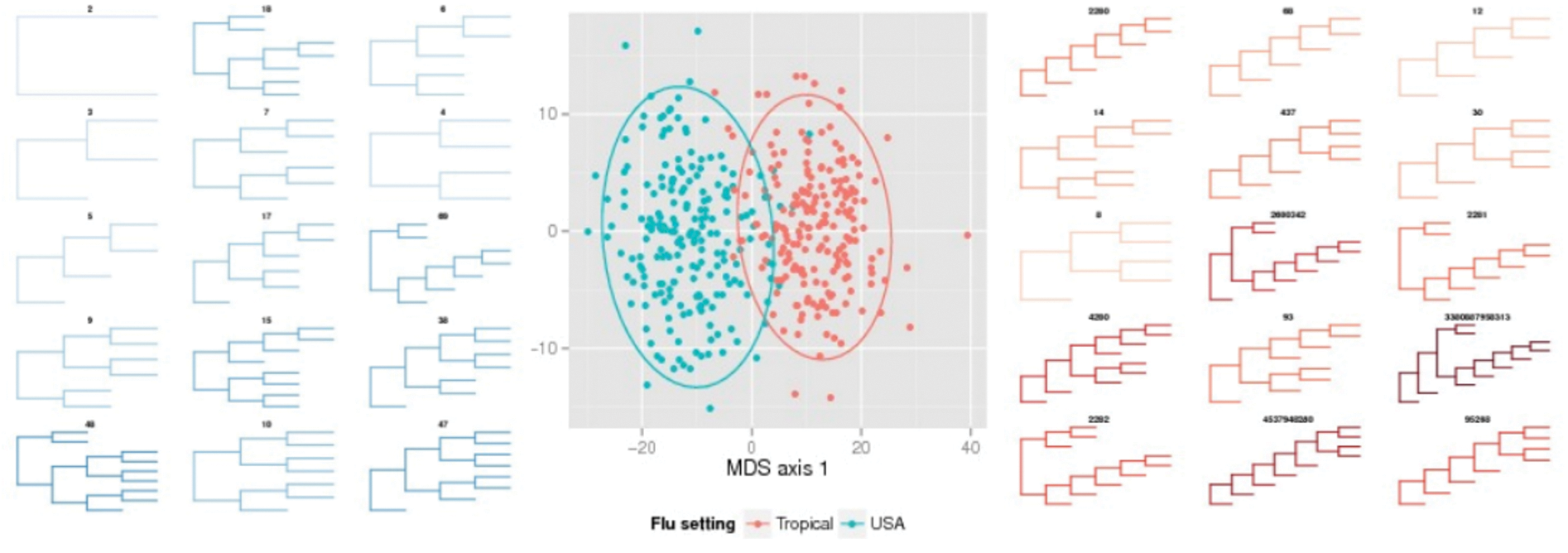
Comparisons between trees from H3N2 flu virus samples. Central panel: multidimensional scaling plot showing that the metric separates trees from the tropics (red) and from the USA (blue). Left and right panels: top-ranked sub-trees that distinguish the two groups, as determined by discriminant analysis of principal components (DAPC); labels correspond to the labelling scheme. Depth of colour corresponds to Sackin imbalance (see Supplementary information).

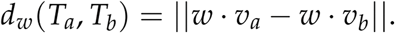

The same weighting can of course be applied to 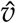 in 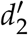.

The labelling scheme induces natural metrics on tree shapes, but does not capture the lengths of branches. These are biologically relevant in many examples, because they reflect the (inferred) amount of time or genetic distance between evolutionary events, although particularly for branches deep in the tree structure they may be difficult to infer accurately. Branch lengths or other features of trees – including the number of lineages through time, diversity measures, tip-tip distance profiles, imbalance measures and bootstrap values – can be incorporated into metrics based on our scheme. As our metric satisfies 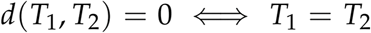, any distance function of the form

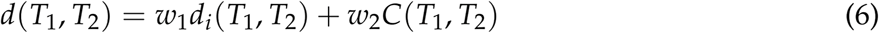

where *C*(*T*_1_, *T*_2_) obeys the triangle inequality will be a metric (though not necessarily Euclidean), even if the features in the comparison *C* do not uniquely define a tree. In Eq. (6), *d* is a metric if *C* is a pseudo-metric.

We can create Euclidean metrics that combine lengths and other features with our shape comparisons. To do this, we describe trees *a* and *b* with vectors *V_a_* and *V_b_*. The first *F* components of *V* capture *F* comparable summaries or length-based statistics, and the remaining components count the label frequencies (as in *v*). In this way we can create any number of Euclidean metrics

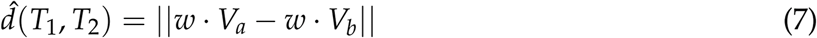

where *w* reflects weightings across the label numbers and summary features. Summary features or comparisons could include spectral differences, Sackin or Colless imbalance, Kullback-Leibler divergence between lineages-through-time plots, maximum likelihood parameter estimates, mean bootstrap values, bootstraps corresponding to each shape label, or other features. Since 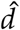 can only be equal to 0 if *T*_1_ and *T*_2_ are the same shape (because of the components of *V* that include the shape label), 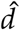 is a Euclidean metric. This extends the shape metric to incorporate branch lengths and to emphasize features of interest (believed to be informative of an underlying process), while retaining the advantages of a true distance metric. Figure S4 illustrates this approach on the tropical and USA trees, showing an MDS plot of 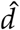, where the first component of *V* is the ratio of the mean terminal branch lengths to the mean internal branch lengths in each tree. While the main shape separation between tropical and USA tree shapes is preserved, there is an informative length dimension illustrating the presence of outliers with high mean terminal branch length.

### 3.5 A convex metric on tree shapes

Mapping tree shapes to other sets of numbers can help us to capture the space of tree shapes in new ways. A particularly nice property of a metric space is convexity - if given two trees *T*_1_ and *T*_2_, there exists a tree *T*_3_ lying directly between them, i.e. 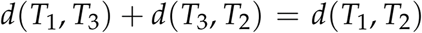. Convex metrics are appealing because in a convex metric on tree shapes we can find the average tree shape for a set of trees, define a centre of mass shape, and further develop statistics on the space of tree shapes.

We use the labelling scheme and a pairing of maps to construct a convex metric on tree shapes. To do this, we map tree shapes to the rational numbers, where the usual absolute value function is a convex metric (as there is always a rational number directly in between any two others). We use the prime decomposition, i.e. the unique product of prime factors of a number (e.g. 10 = 2 · 5). For a tree shape corresponding to integer *n*, we apply 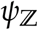 to the exponents of all the prime factors of *n* + 1, and multiply the result (see Methods). For example 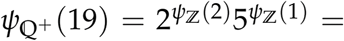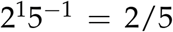. We denote this map 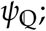 it takes each integer to a unique rational number, and vice versa (bijective). Applying 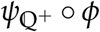 to tree shapes maps them bijectively to the non-negative rational numbers. We can add or multiply trees’ corresponding rational numbers to perform operations in the space of tree shapes. In particular, we can use the usual absolute value distance function to define a convex metric space of tree shapes 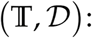

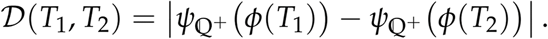

In this space we can find the average tree of a group of trees, and a ‘direct path’ between two trees. Given *n* trees, the average tree is:

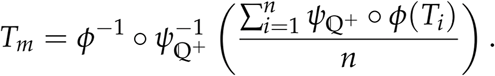

In other words, the average of a set of trees is the tree corresponding to the average of the trees’ rational numbers under the map we have defined. Figure 4 illustrates this operation.

There are infinitely many ways that we could map tree shapes to rational numbers, and we have chosen one that is relatively easy to write down explicitly. Any of them would give rise to a convex metric on the set of tree shapes. It would be most desirable if the resulting metric had some intuitive features - for example, if the trees lying directly between trees *T*_1_ and *T*_2_ (with *n*_1_ and *n*_2_ tips) had an intermediate number of tips between *n*_1_ and *n*_2_ inclusively. The convex metric we have constructed does not have this property, and indeed, since the path between any two rationals traverses a countable infinity of other rationals, but there is a finite number of trees with between *n*_1_ and *n*_2_ tips, no such metric can exist. This convex metric also relies on the prime factorisation of the tree labels, which is a challenge if large labels are encountered.

**Figure 4:**
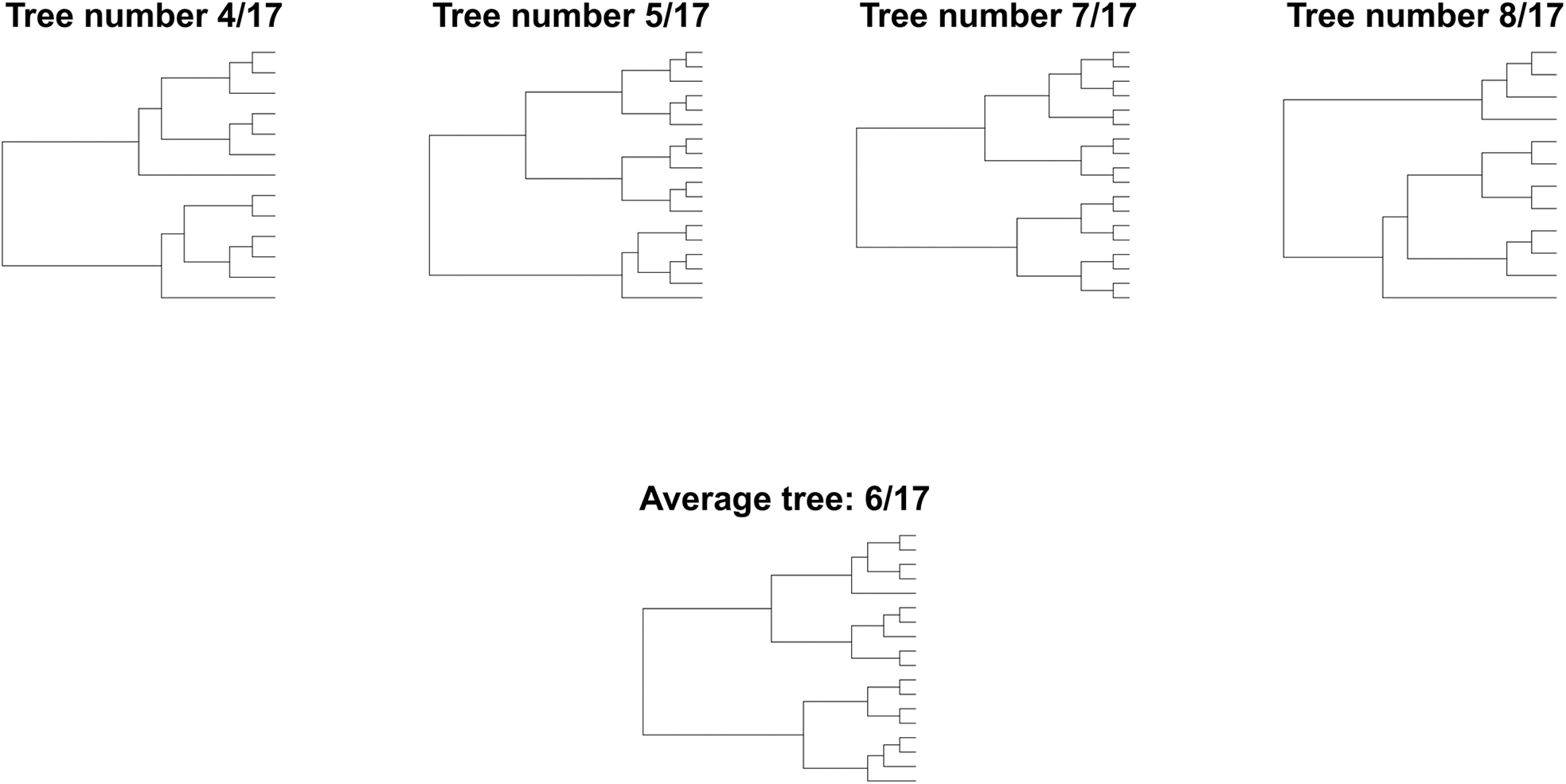
Trees associated with the rationals 4/17, 5/17, 7/17, 8/17, using the map in Example 2. Because the natural distance is convex in 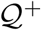, it is possible to find the “average” tree, which is the one mapped into 6/17. Moreover, trees mapped to 5/17, 6/17 and 7/17 are part of the direct path between the trees mapped to 4/17 and 8/17.

## 4 Discussion

We have developed metrics on unlabelled tree shapes, and used them to compare simulated and data-derived trees. The labelling scheme on which the metrics are based comprises a complete characterization of rooted tree shapes, and is not limited to bifurcating trees. Trees from processes known to produce different shapes are well separated in the metric that arises naturally from the scheme. This suggests applications in inferring evolutionary processes and to detecting tree shape bias (Huelsenbeck & Kirkpatrick, 1996; Gascuel, 2000; Stam, 2002). The structure and simplicity of this comparison tool carry a number of advantages. Metrics have good resolution in comparing trees because the distance is only zero if tree shapes are the same. Empirical distributions of sub-tree shapes can easily be found and compared. And as we have shown, the approach can be extended to convex metrics on tree shapes, allowing averaging as well as algebraic operations (addition, multiplication) in tree space. However, this approach does not seem likely to give rise to analytically tractable distributions of tree-tree distances, and in some cases, may not offer more useful resolution than a well-chosen collection of summary statistics.

In particular, scalar measures of asymmetry perform well in distinguishing rooted binary trees. Here, imbalance measures perform slightly worse on the continuous-time birth-death models with *R*0 = 2, 5 but are different between the Yule, PDA, biased and Aldous’ random processes. Matsen (Matsen, 2007) developed a method to define a broad range of tree statistics. Genetic algorithms uncovered tree statistics that can distinguish between the reconstructed trees in TreeBase (Sanderson, Donoghue, Piel, & Eriksson, 1994) and trees from Aldous’ *β*-splitting model, whereas imbalance measures do not (Blum & François, 2006). However, the search-and-optimize approach is vulnerable to over-fitting, as the space of tree statistics is large. It is also reasonable to believe that due to ongoing decreases in the cost of sequencing, studies will increasingly analyze large numbers of sequences and reconstructed trees will have many tips. Any single scalar measure will likely be insufficient to capture enough of the information in these large trees to perform inference.

Large trees present a problem for many approaches to inference, including phylodynamic methods that rely on computationally intensive inference methods. In contrast, our scheme is better able to distinguish between groups of large trees than small ones (fewer than 100 tips). The tip-to-root traversal means that it is very efficient to construct the label set on very large trees (and the same traversal could, with little additional computation time, compute other properties that are naturally computed from tip to root, such as clade sizes, some imbalance measures and many of Matsen’s statistics (Matsen, 2007)). However, due to the large number of tree shapes, the labels themselves become extremely large even for relatively small trees. Our implementation used MD5 hashing to solve this problem, but hashing removes the ability to reconstruct the tree from its label. Also, there are 2^128^ 3 ≈ 10^38^ possible hashed strings, which while large is less than the number of possible tree shapes, even restricting to 500 tips. Alternative labelling schemes may partially alleviate this, for example by subtracting from the label the minimum label for *n* tips, and only comparing trees of size *n* or greater. A related approach was used by Furnas (Furnas, 1984) in developing algorithms to sample trees.

The large size of the labels is also a challenge when they are mapped to 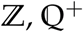 or 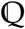 to define a tree algebra or a convex metric. Small changes in the label value can determine visible changes in the shapes. Because the bijective maps are sensitive to small perturbations, the implementation requires the full label, with no hashing compression. However, for trees with 500 tips, we encountered labels of about one million digits. Handling such large numbers with full accuracy required heavy and slow computation. The search for the average tree as found in Figure 4 was only possible for small trees, as the map requires the prime factorization of the label.

Our scheme captures only the shape of the trees; there does not appear to be a natural way to incorporate branch lengths other than appending statistics of branch lengths to the vectors describing the tree (as we have done, though this could be done for each label rather than in aggregate, with a cost for the size of the vector). There are several non-metric approaches to comparing unlabelled trees that do include lengths. In particular, Poon’s kernel method (Poon et al., 2013) compares subset trees that are shared by two input trees, after first "ladderizing" the trees (arranging internal nodes in a left-right order with branching events preferentially to one side). Using a kernel function, this approach can quantify similarity between trees. One challenge is that differences in overall scaling or units of the branch lengths can overwhelm structural differences. Lengths can of coures be re-scaled (for example such that the height of both trees becomes 1), but results may be sensitive to outliers or to the height of the highest tip in the tree. Lengths could also be set to 1 to compare shapes only. Recently, Lewitus and Morlon (LM) (Lewitus & Morlon, 2015) used the spectrum of a matrix of all the node-node distances in the tree to characterise trees; this is naturally invariant to any node and tip labels. They used the Kullback-Leibler divergence between smoothed spectra as a measure of distance. If the spectrum uniquely defined a tree this would be a metric, as it is non-negative and obeys the triangle inequality. As it uses all node-node distances, this approach, requiring the spectrum of a non-sparse 2*n* – 1 × 2*n* 1 matrix for a tree of *n* tips, will become infeasible for large trees.

One option is to add one or several terms to the distance function to incorporate more information, as outlined above. Combinations of our distances and other tree comparisons may turn out to be the most powerful approach to comparing unlabelled trees, allowing the user to choose the relative importance of scalar summaries, tree shape, spectra and so on while retaining the discriminating power of a metric. Ultimately, discriminating and informative tools for comparing trees will be essential for inferring the driving processes shaping evolutionary data.

